# Controlled experiment finds no detectable citation bump from Twitter promotion

**DOI:** 10.1101/2023.09.17.558161

**Authors:** Trevor A. Branch, Isabelle M. Cȏté, Solomon R. David, Joshua A. Drew, Michelle LaRue, Melissa C. Márquez, E. Chris M. Parsons, D. Rabaiotti, David Shiffman, David A. Steen, Alexander L. Wild

## Abstract

Multiple studies across a variety of scientific disciplines have shown that the number of times that a paper is shared on Twitter (now called X) is correlated with the number of citations that paper receives. However, these studies were not designed to answer whether tweeting about scientific papers causes an increase in citations, or whether they were simply highlighting that some papers have higher relevance, importance or quality and are therefore both tweeted about more and cited more. The authors of this study are leading science communicators on Twitter from several life science disciplines, with substantially higher follower counts than the average scientist, making us uniquely placed to address this question. We conducted a three-year-long controlled experiment, randomly selecting five articles published in the same month and journal, and randomly tweeting one while retaining the others as controls. This process was repeated for 10 articles from each of 11 journals, recording Altmetric scores, number of tweets, and citation counts before and after tweeting. Randomization tests revealed that tweeted articles were downloaded 2.6–3.9 times more often than controls immediately after tweeting, and retained significantly higher Altmetric scores (+81%) and number of tweets (+105%) three years after tweeting. However, while some tweeted papers were cited more than their respective control papers published in the same journal and month, the overall increase in citation counts after three years (+7% for Web of Science and +12% for Google Scholar) was not statistically significant (*p* > 0.15). Therefore while discussing science on social media has many professional and societal benefits (and has been a lot of fun), increasing the citation rate of a scientist’s papers is likely not among them.

## Introduction

Scientists are increasingly encouraged to communicate their science and “escape from the ivory tower”, engaging with the public, the press, and policymakers [1]. It is argued that scientists need to be part of a social contract, devoting their energy to the most pressing problems of our time and communicating and explaining their findings [2]. Many scientists have taken this advice to heart and devoted considerable time to science communication using social media platforms, and in turn a cottage industry of alternative metrics has arisen that measures the impact of scientific papers beyond simple citation counts [3-5]. Key among these is the Altmetric attention score from altmetric.com, which combines attention from newspapers, blogs, and social media to collectively formulate a single score for individual scholarly papers [6]. Despite the broad sweep of Altmetric.com, the resulting scores are usually dominated by counts of the number of tweets posted on Twitter [7-9]. We note in passing that Twitter was recently renamed “X”, and tweets to “reposts”, but since even AP style guides suggest “X, formerly known as Twitter”, indicating that the brand change is not widely recognized yet, we continue to refer to Twitter and tweets here.

A longstanding discussion in the world of science communication is this: does science communication—in addition to educating broadly—have an additional benefit of increasing the profile of scientific papers, resulting in a higher number of citations? In almost every scientific field examined, ranging from ornithology to urology, there are significant correlations between Altmetric scores (or number of tweets) and the number of citations that a paper receives [7-18], but correlation does not prove causation. An alternative explanation, termed “good papers are good” (C. McClain, pers. comm.) posits that research findings that are exciting, novel, and timely, and that are of substantial interest to the wider public, will both receive a lot of online attention and be cited widely; and thus correlation between tweets and citations does not prove that tweets cause more citations. Proof can come only in the form of a controlled scientific experiment where individual articles are randomly assigned to be in the experimental arm (i.e., promoted on social media) or the control arm (i.e., not promoted). After some discussion about this idea on Twitter in 2018 (where else!), the authors of this study conceived and initiated a controlled experiment to examine the impact of tweeting papers on citation counts.

Previous controlled experiments to investigate the impact of social media on citations have resulted in contradictory findings [19-24]. The earliest effort randomized 130 papers from the *International Journal of Public Health*, promoting half on the journal’s blog, Facebook, and Twitter [24, 25]. After 24 months, there was no significant impact on downloads (428 vs. 423, *p* = 0.84) or citations (4.11 vs. 3.65, *p* = 0.70). However this journal has a relatively low citation rate, and the associated Facebook and Twitter accounts had low follower numbers at the onset (140 and 403 respectively) [24]. A larger-scale experiment, the ESC Journals Randomized Study [20, 21], randomly picked 694 articles, promoting half of them on Twitter, and found 1.12 times more citations (95% CI 1.08–1.15, *p* < 0.0001) among the tweeted articles. However, the Twitter promotion was also associated with 24 hours of free access to the articles [20, 21], which may have increased citations. The analysis also ignored overdispersion, which if accounted for, would have resulted in a non-significant (*p* = 0.17) outcome [26]. The next effort involved an intensive randomized trial, where four articles per day were randomly selected and tweeted by 13 accounts (with combined 52,983 followers). The authors reported that citations increased far more for the tweeted than the control articles (3.1 vs. 0.7) after one year [22, 27], but proved controversial, with an outside reanalysis finding no citation benefit [28], although the authors stand by their study [29]. One final experiment involved a promotion campaign on 24 experimental vs. 24 control papers in *Cephalalgia* in 2019-2020, including press releases, custom graphics, tweets, and promotions on Facebook, Reddit, Youtube, Wikipedia, news outlets, blogs, and more [23]. After 24 months, the Altmetric score of the promoted papers was far higher (55.6 vs. 8.1), while the number of citations increased by 17% on Dimensions Citations (*p* = 0.04), by 19% on CrossRef (*p* = 0.04), and by 11% on Web of Science (but this increase was not significant, *p* = 0.18).

The above experiments were part of intensive marketing and promotion campaigns by journals or associations [23-25], included free article access to promoted papers [20, 21], or were tweeted by multiple Twitter accounts [22]. The very fact that a paper is being heavily promoted might signal to readers that the paper is worth downloading (and later citing), but this is atypical for the vast majority of science Twitter use, which consists of individual scientists highlighting a scientific paper with one or two tweets. Our experiment therefore set out from the onset to test whether scientists with high follower counts posting naturally on Twitter can increase citations of scientific papers, expecting that tweeted articles would be associated with higher Altmetric scores, and receive more citations over a three-year period than control articles for each journal.

## Materials and methods

The experiment was intended to mimic the natural tweeting patterns of each participant, to address whether scientists with large social media followings can increase future citations. The 11 selected tweeters (the coauthors of this study) were all active on Twitter at the time of the experiment and most have sufficient followers (>5,000) to be considered “influential” [30, 31]. The authors also all have a long-standing interest in using social media for science communication. The tweeters were followed by a mean of 15,257 (range 3,962–39,429) accounts at the start of the tweeting experiment and 20,310 (4,647–47,842) at their final experimental tweet; eight have >10,000 followers; and four have >20,000 followers. For context, a list of 307 university faculty on Twitter in aquatic and fishery sciences (compiled by T.A.B.) (https://twitter.com/i/lists/105667923) included only 4.6% with > 10,000 followers (compared to 73% of our authors).

### Experimental design

Each of the 11 participants selected one journal in their field on a first-come-first-served basis, with the intent for each to tweet one randomly chosen article per month for 10 months, for a total of 110 tweeted articles (Table 1). Each month of the experiment, the coordinator (T.A.B.) identified all eligible papers from online-early articles in the respective journals published in the most recent 30 days, or from the latest online issue, depending on the journal. Articles had to be primary research articles or reviews, and we excluded short notes, book reviews, corrections, letters to the editor, opinion pieces, and other non-standard types of articles. We also excluded articles that were already being highlighted by the journal (e.g., editor selected papers), coauthored by any of the participants in the study, or represented a conflict of interest to the tweeter. Occasionally, a waiting period was required to ensure at least five articles met the criteria, thus no individual account tweeted about more than one paper per month. From all eligible articles, a random number generator was used to select five, with the first selection becoming the experiment (tweeted), and the other four becoming the controls. We chose four controls instead of one since this increases the statistical power of the experiment at little extra cost.

**Table 1:** Journal and tweeter summary. List of journals sorted by 2021 impact factor (Web of Science), the account that tweeted articles from that journal, with follower numbers at first and last tweet; the mean number of link clicks 30 days after tweeting (i.e., clicks sending them to the journal article); and mean Twitter impressions 30 days after tweeting (number of views of the tweet). Each account tweeted 10 articles from each journal.

| Journal name | Impact factor | Twitter handle | Followers start | Followers end | Link clicks | Tweet impressions |
| --- | --- | --- | --- | --- | --- | --- |
| Conservation Biology | 7.56 | @Drew_Lab | 7,139 | 8,402 | 13.7 | 2,650 |
| Journal of Animal Ecology | 5.61 | @DaniRabaiotti | 10,878 | 19,611 | 37.1 | 4,465 |
| Coral Reefs | 4.64 | @redlipblenny | 6,236 | 7,348 | 21.8 | 2,722 |
| Marine Policy | 4.32 | @WhySharksMatter | 39,429 | 47,842 | 2.3 | 1,715 |
| ICES Journal of Marine Science | 3.91 | @TrevorABranch | 9,909 | 12,120 | 54.2 | 5,106 |
| Marine Ecology Progress Series | 2.92 | @mcmsharksxx | 11,913 | 17,322 | 3.8 | 1,868 |
| Journal of Wildlife Management | 2.59 | @AlongsideWild | 31,281 | 42,330 | 17.7 | 4,051 |
| Ecological Entomology | 2.23 | @myrmecos | 20,938 | 25,236 | 27.5 | 4,935 |
| Transactions of the American Fisheries Society | 2.20 | @SolomonRDavid | 11,483 | 16,943 | 16.4 | 2,358 |
| Polar Biology | 2.20 | @drmichellelarue | 14,655 | 21,609 | 5.9 | 2,407 |
| Journal of Wildlife Diseases | 1.63 | @Craken_MacCraic | 3,962 | 4,647 | 0.6 | 429 |
| Overall mean | 3.62 |  | 15,257 | 20,310 | 18.3 | 2,973 |

The participants were informed of the five articles (one to tweet, the other four to avoid tweeting) and instructed to post a tweet as soon as possible that summarized the main message of the paper in the same manner as they would usually tweet about scientific papers, for example including a brief summary of the highlights of the paper or a quote from the abstract or conclusions. They were additionally instructed to include a link to the paper itself, and an image, figure, or other illustration (e.g. Fig. 1). They were instructed to engage with any replies in their normal manner. Tweeters signed up for Twitter Analytics (www.analytics.twitter.com) to obtain information about tweet impressions after 30 days.

**Fig. 1.**
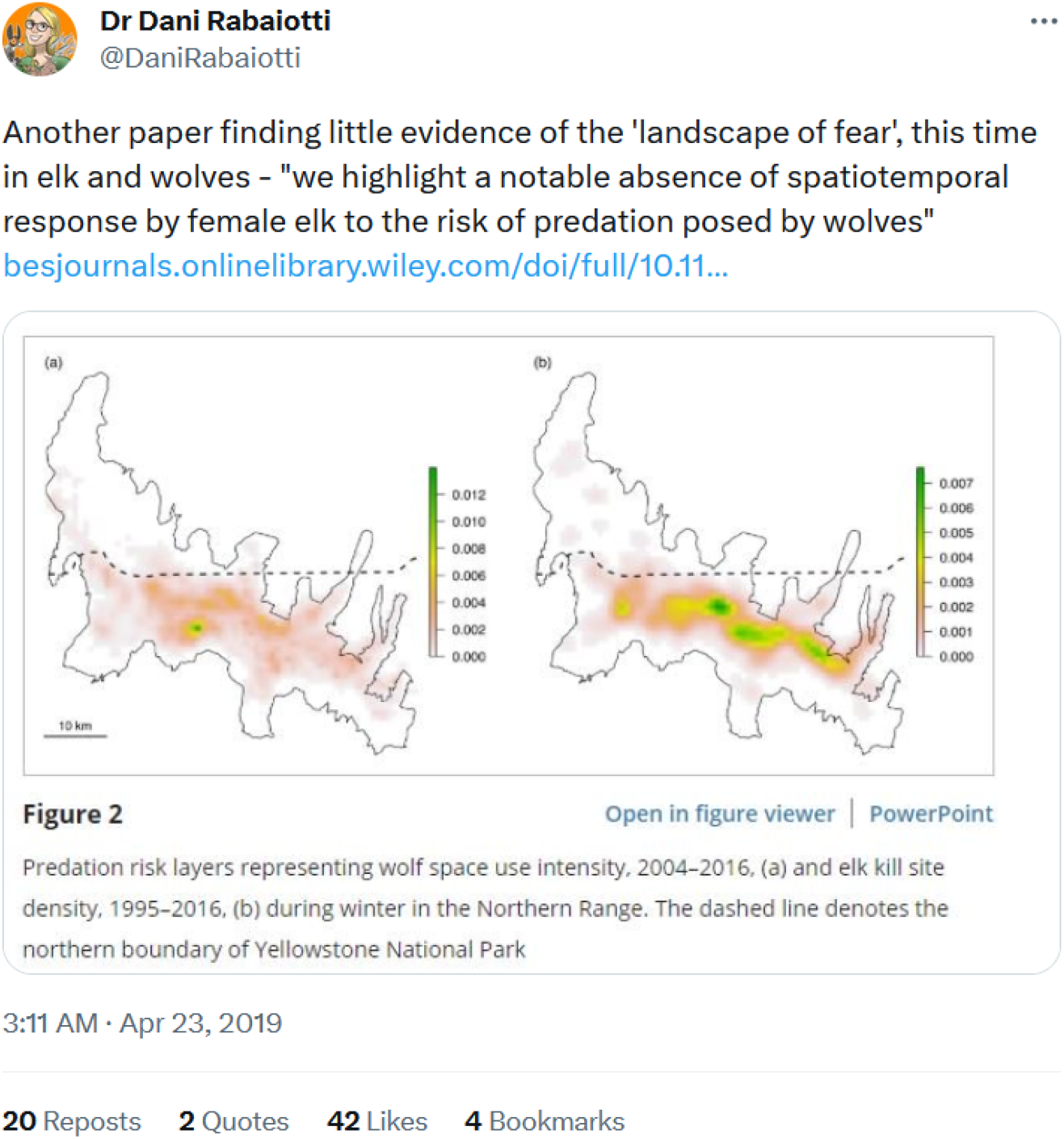
**Example experimental tweet**, including a short description of the key finding of the paper, a link to the paper, and one of the figures in the paper. Within 30 days, this tweet received 10,032 view impressions, and 99 link clicks sending viewers to the linked paper.

The first period of data collection occurred just before each tweet: the coordinator recorded the current date, number of followers for the tweeter, Altmetric Attention Score from the bookmarklet from altmetric.com, and number of tweeters for each paper, and then alerted the tweeter with links to the selected articles, highlighting which article to tweet. If there was a delay in tweeting, the “before” metrics were updated before sending a reminder email. Citation counts on Google Scholar and Web of Science were zero for nearly every paper in the first few months of the experiment and were therefore not systematically collected. Due to holiday periods and delays in tweeting, the entire experiment took longer than the intended 10 months, encompassing 5 December 2018 to 16 April 2020.

The second period of data collection was 30 days after each tweet. Social media attention metrics are highest soon after publication, and then dwindle rapidly [12, 31, 32]. At 30 days, the coordinator collected the Altmetric attention score and number of tweeters for all five papers, and asked the tweeter to obtain Twitter analytics for their tweet to get the number of tweet impressions and link clicks to the paper in the experimental arm (Table 1).

The final period of data collection was three years after each tweet, with final data collection ending on 16 April 2023, and consisted of Altmetric score, number of tweeters, number of citations from Google Scholar, and number of citations from Web of Science. Throughout this process, participants were instructed not to tell their followers that an experiment was underway so that the impact of the tweets would be as close as possible to their natural tweeting patterns of occasionally highlighting scientific papers.

### Human subjects approval

The need for consent from users of Twitter was waived by the University of Washington Institutional Review Board.

### Impact on daily downloads

To assess the immediate impact of tweets on article downloads, we obtained daily download counts for articles in five of the journals (all published by John Wiley & Sons): *Conservation Biology, Journal of Wildlife Management, Journal of Animal Ecology, Ecological Entomology*, and *Transactions of the American Fisheries Society*. Data were obtained for each day for two years after the likely publication date. For each group of five articles, we found the downloads on the day of tweeting (day 0) and each day thereafter (day 1, 2, 3, etc.), and compared the corresponding daily mean download counts. We could not compare time series of downloads before the tweet date because some articles were tweeted on the likely publication date. (For one control article with a likely publication date one day *after* the tweet date, we subtracted one day so that the likely publication date equaled the tweet date.)

### Improvements over previous experiments

The experimental design that we used offers several improvements over previous experiments. Notably, the power to detect a change is increased by tracking four control papers for every tweeted paper, instead of one control for each tweeted paper. We also controlled for citations accumulating over time by fixing the data collection period at 3 years after the tweeting intervention, rather than recording data on a fixed date with differing amounts of time since the intervention. Most importantly, the intervention is designed to mimic the natural process of encountering a paper and tweeting about it, to see whether this typical intervention has an impact on the tweeted paper. Other experiments have involved mass tweeting from multiple accounts on the same day, multiple papers being tweeted within a limited amount of time (a few days to weeks), or mass marketing campaigns across multiple forms of media [22, 23]. In our experiment, each individual tweeted about an assigned paper included in the experiment no more than once per month, so that it would not be obvious to readers that a deliberate experiment was taking place. Finally, we used high-follower accounts to do the tweeting. Should the intervention from highly influential accounts have no impact on citations, we can conclude that smaller accounts would likely also have little impact.

### Analysis

We used randomization tests for significance for each of these metrics: Altmetric scores 30 days and 3 years after tweeting, number of tweeters after 30 days and 3 years, number of citations in Web of Science after 3 years, and number of citations in Google Scholar after 3 years. The randomization tests were conducted as follows (using as an example Google Scholar citations after 3 years). We calculated the test statistic (e.g., 1.4 citations) as the difference in mean citations between the tweeted (e.g., 13.4) and control (e.g., 12.0) articles. To test the likelihood that this difference could have arisen by chance alone, we randomly reassigned the label “tweeted” and “control” for the 110 sets of articles: within each set of five articles, we randomly chose one article (and labeled it “tweeted”), and then labeled the other four as “controls”, then repeated this process for all 110 sets of 5 articles, and calculated *X*_1_ = mean(“tweeted”) – mean(“control”). We repeated this sampling process 100,000 times to obtain the sampling distribution (*X*_1_, *X*_2_, …, *X*_100000_). The *p*-value is the proportion of times that values in the sampling distribution are greater than the test statistic. Similar tests were conducted for each article-level metric.

In addition to the randomization tests based on the raw counts for each article, we also conducted tests based on normalized citation counts, to ensure that each journal contributed equally to the tests rather than journals with high citation counts contributing more. Normalizing involved calculating the mean and SD of citations for all 50 articles in a journal, and then converting the counts for each article by as follows: [citation – mean(citations)] / SD(citations). Randomization tests (as outlined in the previous paragraph) were conducted on the normalized values.

## Results

We tweeted a total of 110 papers, each matched with four control papers that we did not tweet, for a total of 550 tracked papers. The dataset we compiled is posted under Supplementary Materials for transparency. During the experiment, the tweeting accounts continued to grow in influence (mean

5,053 followers added) (Table 1). Each tweet garnered a mean of 2,973 impressions and 18 link clicks to each scientific paper (Table 1).

Daily downloads (for five journals) averaged 3.9 times higher on the day of tweeting for the tweeted articles compared to the controls (range 2.4–6.9, Fig. 2, Table 2). After one day, daily downloads were still higher for tweeted articles in all journals (mean 2.6, range 1.3–5.1 times), but thereafter not all journals had higher downloads for tweeted articles (Table 2). Moreover, overlapping intervals of ±1 SE suggested a diminished effect (Fig. 2). Since some tweets were sent out on the same day articles were published online, a full comparison was not possible for the days preceding the tweets, but there did appear to be download increases for tweeted articles in some journals on the day immediately before tweeting occurred, likely attributable to time zone differences in the tweets vs. the database of downloads.

**Table 2.** Ratio of daily download counts for tweeted articles vs. control articles. Geometric means (and not arithmetic means) are calculated across journals since the metrics are ratios.

| Journal name | Day 0 | Day 1 | Day 2 | Day 3 | Day 4 | Day 5 |
| --- | --- | --- | --- | --- | --- | --- |
| Conservation Biology | 3.3 | 1.3 | 1.2 | 1.8 | 1.0 | 1.2 |
| Ecological Entomology | 3.0 | 4.0 | 3.2 | 1.5 | 1.7 | 7.7 |
| Journal of Animal Ecology | 5.5 | 5.1 | 2.5 | 1.4 | 1.2 | 2.1 |
| The Journal of Wildlife Management | 6.9 | 2.1 | 2.4 | 1.4 | 0.2 | 1.8 |
| Transactions of the American Fisheries Society | 2.4 | 2.2 | 0.4 | 4.7 | 8.0 | 0.8 |
| Geometric mean | 3.9 | 2.6 | 1.6 | 1.9 | 1.3 | 2.0 |

**Fig. 2.**
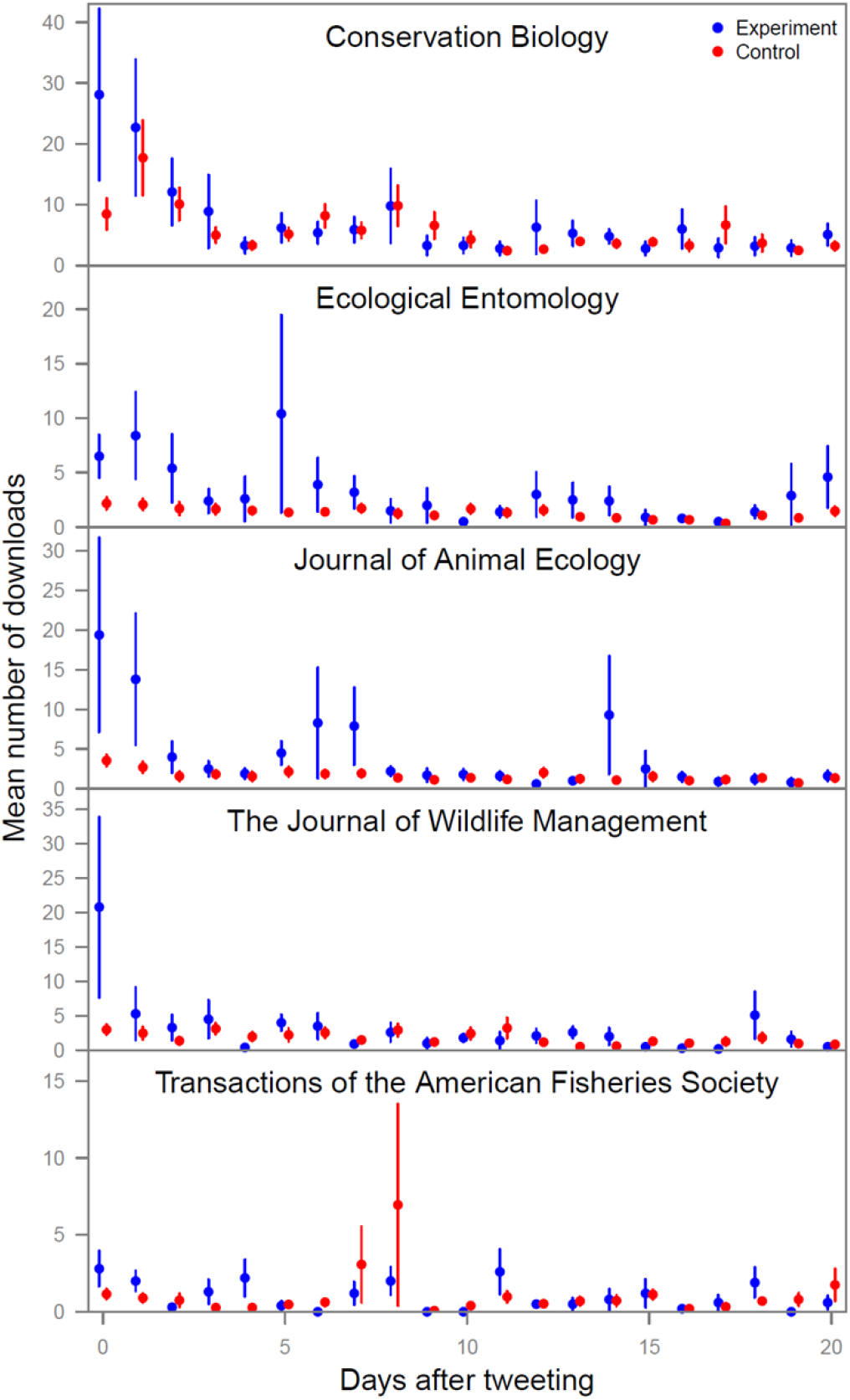
Daily article downloads for tweeted vs. control articles after the tweeting date. Data obtained from John Wiley & Sons for five journals for the experimental (blue, *n* = 10 per journal) and control (red, *n* = 40 per journal) articles, aligned to the number of days after tweeting occurred for each experimental article. Points are means, lines are ±1 SE.

After 30 days, accumulated Altmetric scores for tweeted articles were 68% higher than the controls (*p* = 0.065), and the number of tweets was 64% higher (*p* = 0.004) (Table 3). When articles were normalized to ensure that each journal counted equally, differences for Altmetric scores and number of tweets were highly significant (*p* < 0.0001, Table 3). It should be noted that the increase in Altmetric score (by 7.3) includes 0.5 from the experimental tweet, and the increase in number of tweets (by 6.8) includes 1.0 from the experimental tweet. Similar patterns were seen 3 years after tweeting: the Altmetric score was 81% higher than controls (*p =* 0.004) and the number of tweets was 105% higher (*p* = 0.004), with lower *p*-values (<0.0001) when normalized (Table 3).

**Table 3.** Comparison of article metrics between the tweeted and control articles. Mean Altmetric score and number of tweets about the articles both 30 days and 3 years after tweeting (for all 11 journals), and mean number of citations in Web of Science and Google Scholar three years after tweeting. The p-values were obtained by resampling—either the observed counts (“p”)—or after normalizing by subtracting the mean and dividing by the standard deviation for the respective journals (“p normalized”).

| Metric | Elapsed time | Mean tweeted | Mean control | Tweeted/control | Tweeted – control | $p$ | $p$ normalized |
| --- | --- | --- | --- | --- | --- | --- | --- |
| Altmetric score | 30 days | 18.0 | 10.7 | 1.68 | 7.3 | 0.065 | <0.0001 |
| Altmetric score | 3 years | 23.0 | 12.7 | 1.81 | 10.3 | 0.024 | <0.0001 |
| Number of tweets | 30 days | 17.4 | 10.7 | 1.64 | 6.8 | 0.004 | <0.0001 |
| Number of tweets | 3 years | 24.0 | 11.7 | 2.05 | 12.3 | 0.004 | <0.0001 |
| Web of Science citations | 3 years | 8.6 | 8.0 | 1.07 | 0.6 | 0.258 | 0.709 |
| Google Scholar citations | 3 years | 13.4 | 12.0 | 1.12 | 1.4 | 0.157 | 0.415 |

We expected that Altmetric scores would be similar after 30 days and after 3 years, and indeed the Altmetric scores after 30 days were 78% of the 3-year score for tweeted articles, and 84% for control articles. The number of tweets after 30 days was 73% of the 3-year score for tweeted articles but considerable higher (91%) for control articles. This difference (73% vs. 91%) may be due to chance or might hint at a more long-lasting effect beyond 30 days if articles are tweeted by “influential” accounts.

The number of citations after 3 years was higher for tweeted articles than controls, but these differences were not statistically significant: for Web of Science, citations increased by 0.6 (7%) per article (*p* = 0.258), and for Google Scholar by 1.4 (12%) per article (*p* = 0.157). Normalizing increased the *p*-values (to 0.709 and 0.415 respectively).

It is possible that our experiment did not have enough statistical power to detect an increase in citations. To estimate the sample size required to detect a true effect size of 7% (Web of Science) and 12% (Google Scholar), we replicated the entire dataset 2, 3, 4, …, *n* times to find the sample size at which *p* < 0.05 from the resampling analysis (with 10,000 simulations). We found that ≥ 770 tweeted articles would be needed to detect a true difference of this size in Web of Science citations, and ≥ 330 tweeted articles for Google Scholar (Table 4), which are both far higher than in our experiment (*n* = 110).

**Table 4.** Power to detect differences in citations of tweeted articles. Estimated number of tweets required to have found a significant difference in citations (*p* < 0.05), if the observed effect is the true effect size. This was calculated by replicating the data 2–10 times, repeating the resampling analysis, and finding the smallest sample size with *p <* 0.05 (in bold) for Web of Science and Google Scholar.

| Data<br>duplicates | $n$<br>tweeted | p(Web of<br>Science) | p(Google<br>Scholar) |
| --- | --- | --- | --- |
| 1 | 110 | 0.257 | 0.155 |
| 2 | 220 | 0.187 | 0.082 |
| 3 | 330 | 0.133 | **0.039** |
| 4 | 440 | 0.104 | 0.023 |
| 5 | 550 | 0.074 | 0.010 |
| 6 | 660 | 0.057 | 0.007 |
| 7 | 770 | **0.047** | 0.004 |
| 8 | 880 | 0.035 | 0.002 |
| 9 | 990 | 0.026 | 0.002 |
| 10 | 1100 | 0.023 | 0.001 |

## Discussion

In this experiment, tweeting about a scientific paper resulted in an average of 2,973 impressions and 18 link clicks to the article, in addition to a 3.9-fold increase in downloads of the article on the day of tweeting, and 2.6-fold increase the day after tweeting. Furthermore, within 30 days, Altmetric scores were 68% higher than for control articles, due largely to a 64% increase in the number of tweets about the article; and this difference continued to increase so that after three years, the Altmetric score was 81% higher and the number of tweets 105% higher. Thus, by the broader measure of alternative metrics, tweeting by Twitter-influential scientists raised the profile of the tweeted articles compared to the controls. In other words, more people (including scientists and non-scientists) became aware of, downloaded, and possibly even read these papers than would have otherwise.

However, tweeting did not result in significantly higher citation counts—one perceived metric of the overall quality and importance of a scientific paper—within three years. Three years is generally sufficient for citations of articles to approach asymptotic annual values [e.g., 33]. Although citations for tweeted articles were 7% higher in Web of Science, and 12% higher in Google Scholar, these differences were not statistically significant whether based on raw counts or after normalizing to ensure that all journals counted equally. If these are close to the true effect sizes, our experiment did not have sufficient power: we would need 3 times (Google Scholar) to 7 times (Web of Science) greater sample sizes to detect these differences at *p* < 0.05. The average scientist should therefore not expect a detectable increase in citations resulting from tweeting about their papers, especially since the accounts involved in this study have much larger than average follower counts.

A corollary to this point is that the scientific literature itself (unlike Altmetrics) appears to be resilient to attention gaming on social media: more eyeballs on papers does not necessarily result in higher citation counts. Instead, higher citation counts of highly tweeted papers reflect the underlying value of important papers being recognized both by scientists and by social media users (i.e., “good papers are good”).

It should be pointed out that Twitter accounts with large numbers of followers also have a more diverse set of followers [34]: accounts with >10,000 followers typically have as many followers that are members of the general public as they have followers who are scientists. For science communicators, this is a key target audience beyond the ivory tower, but of course members of the public are highly unlikely to publish papers that cite the papers mentioned in tweets. The fact that more members of the public become aware of current research findings from newly published papers has a variety of societal benefits, including scientific literacy, which is a fundamental component of a democratic society [35], just not increased citations of those papers.

Previous controlled experiments either found no effect on citations [24, 25], found a similar sized increase (12%) [21], or found a very large increase in citation counts (+3.1 for tweeted vs. +0.7 for controls) after one year [22], but the experiments finding increases have been criticized [26, 28]. Here we were unable to detect a significant increase in citations despite tweets coming from influential science Twitter accounts with 4,000 to 48,000 followers each.

For more than a decade, Twitter has been an incredibly popular platform for researchers to share their scientific advancements with a broader audience, so it is with a measure of wistfulness that we acknowledge the decline of Twitter in recent months after its purchase, job cuts, and rebranding to X. For example, a survey of nearly 9,200 scientists active on Twitter revealed that 54% have quit Twitter altogether or reduced their Twitter use in the past six months [36], while 46% have opened an alternative microblogging account (the top three recipients being Mastodon, Instagram or Threads) [36].

With the decline of Twitter, we are concerned that there may be a decrease in the rapid dissemination of research, an impediment to cross-disciplinary collaboration and knowledge exchange, and a decline in the ability of scientists to reach and educate a wider audience, thus impacting public understanding of and support for science. The departure of scientists from the platform formerly known as Twitter may in turn leave the platform more vulnerable to the proliferation of misinformation that contradicts accepted scientific findings. Our hope is that any mass departure of scientists from Twitter is matched with a mass entry of scientists to alternative forms of social media providing new opportunities for scientific communication and engagement.

As a group of authors, we have all benefited greatly from our social media foray, building an online community where we learn from others, are filled with wonder at the marvels of nature, and at outrage at the sins of humankind. And some of this work has resulted in scientific papers and collaborations that would not otherwise have happened [37-39], including this very paper. Increasing the profile of our scientific papers was certainly not our primary aim, and thus perhaps the real value of online public science engagement is how many friends we added along the way, and the knowledge we shared with, and gained from, our online communities… not higher citation counts.

## Acknowledgments

No funding was received for this work, which grew from early discussions that included C. R. McClain. We thank the Market and Publishing Analytics Team of John Wiley & Sons for writing scripts to extract daily download numbers for the five journals analyzed here, without which we could not have measured the impact of tweets on paper downloads. We would like to thank the scientists on Twitter who engaged with the tweets in this paper with the enthusiasm and openness that was always the hallmark of the science Twitter community.

